# Self-regulation of the Dopaminergic Reward Circuit in Cocaine Users with Mental Imagery and Neurofeedback

**DOI:** 10.1101/321166

**Authors:** Matthias Kirschner, Ronald Sladky, Amelie Haugg, Philipp Stäempfli, Elisabeth Jehli, Martina Hodel, Etna Engeli, Sarah Höesli, Markus R. Baumgartner, James Sulzer, Quentin J.M. Huys, Erich Seifritz, Boris B. Quednow, Frank Scharnowski, Marcus Herdener

## Abstract

**Background:** Enhanced drug-related reward sensitivity accompanied by impaired sensitivity to non-drug related rewards in the mesolimbic dopamine system are thought to underlie the broad motivational deficits and dysfunctional decision-making frequently observed in cocaine use disorder (CUD). Effective approaches to modify this imbalance and reinstate non-drug reward responsiveness are urgently needed. Here we examine whether cocaine users (CU) can use mental imagery of non-drug rewards to self-regulate the ventral tegmental area and substantia nigra (VTA/SN). We expected that compulsive and obsessive thoughts about cocaine consumption would hamper the ability to self-regulate the VTA/SN. Finally, we tested if self-regulation of the VTA/SN can be improved with real-time fMRI (rtfMRI) neurofeedback (NFB).

**Methods**: Twenty-two CU and 28 healthy controls (HC) were asked to voluntarily up-regulate VTA/SN activity with rewarding non-drug imagery alone, or combined with rtfMRI NFB of VTA/SN activity. Obsessive-compulsive drug use was measured with the Obsessive Compulsive Cocaine Use Scale (OCCUS).

**Results:** CU were able to induce activity in the dopaminergic midbrain and other reward regions with reward imagery. The ability to self-regulate the VTA/SN was reduced in those with more severe obsessive-compulsive drug use. NFB enhanced the effect of non-drug imagery.

**Conclusion:** CU can voluntary activate their reward system through non-drug related imagery and improve this ability with rtfMRI NFB. Combining reward imagery and rtFMRI NFB has great potential for modifying the maladapted reward sensitivity and reinstating non-drug reward responsiveness. This motivates further work to examine the therapeutic potential of cognitive neurostimulation in CUD.

## Introduction

Cocaine addiction is a severe and often chronically relapsing-remitting disorder characterized by loss of control, impulsive and compulsive drug intake often driven by obsessive thoughts about drug use (1, 2). In the transition from recreational substance use to addiction, neuroplastic adaptations within the mesolimbic dopamine system contribute to complex alterations in reward processing (2, 3). In particular, both an enhanced mesolimbic sensitivity to drug-related reward signals, and a reduced sensitivity to non-drug related rewards contribute to dysfunctional decision making and the characteristic narrowing of interests (4, 5). Thoughts obsessively circle around cocaine use, while drug seeking and consumption compulsively dominate behavior at the expense of previously rewarding ones such as social activities or hobbies (6, 7). The clinical relevance
 of this maladaptation has been recognized by forthcoming diagnostic systems (ICD-11), in which imbalanced reward sensitivity will be one of the three defining characteristics of substance dependence (6). At the neural level this maladaptation manifests in increased activity in reward regions like the ventral tegmental area and substantia nigra (VTA/SN) in response to drug-related cues (8–11) and impaired sensitivity in these regions to non-drug rewards like external monetary or social reward cues (12–16). While conventional therapeutic approaches mostly focus on reducing sensitivity to drug-related stimuli, it is unknown how we can modify this imbalance by reinstating non-drug reward responsiveness.

Recent evidence suggests that reward-related neural activation can be self-regulated using feedback of circumscribed brain activity measured online with functional magnetic resonance imaging (fMRI), a procedure known as real-time fMRI neurofeedback (rtfMRI NFB) (17). For example, Sulzer et al. demonstrated that healthy individuals can use reward imagery to self-regulate activation in the ventral tegmental area and substantia nigra (VTA/SN) and that this ability improves with online visual feedback of VTA/SN activity (18). This self-regulation ability was corroborated by two other studies, one focused on VTA (19), and the other on nucleus accumbens (20). While the potential implications of self-regulated reward activity are manifold, its clinical relevance has yet to be realized. Combining reward imagery and NFB, this novel approach allows us to modify reward sensitivity with personalized non-drug rewarding stimuli in individuals with cocaine use disorders (CUD).

The first aim of this study was to probe whether cocaine users (CU) can use non-drug related rewards to endogenously regulate the VTA/SN activity. As sensitivity to non-drug related rewards is thought to diminish during the transition to chronic cocaine use, the ability to gain self-control of reward-related brain regions via non-drug reward imagery might be impaired in individuals with more severe compulsive and obsessive thoughts about cocaine use (21). Therefore, we hypothesize that the severity of compulsive and obsessive thoughts correlates negatively with the VTA/SN activation during mental imagery. The second aim of the study was to investigate whether CU can use rtfMRI NFB to improve the ability to self-regulate the VTA/SN. In summary, we aimed to investigate whether self-regulation of the putatively dopaminergic mesolimbic rewards system with reward imagery is impaired and if NFB might be a suitable approach to improve reduced non-drug reward sensitivity in CUD.

## Methods

### Participants

We recruited 30 CU and 30 healthy controls (HC) were recruited from inpatient and outpatient units of the Psychiatric University Hospital Zurich and via online advertisement. Inclusion criteria for CU were cocaine use of at least 0.5 g/week, cocaine as the primary used illegal drug and current abstinence duration of no longer than 6 months. Self-reports were controlled by urine toxicology and 6-month hair analysis (22, 23). Exclusion criteria for the CU were use of opioids and a polysubstance use pattern other than recreational use. Because of their high prevalence in CU, nicotine dependence, attention deficit hyperactivity disorder and history of depression were not excluded. Other lifetime or current axis I DSM-IV disorders (24) led to exclusion. HC and CU were matched for sex, age and for nicotine consumption. Exclusion criteria for HC were any axis I DSM-IV psychiatric disorder with the exception of nicotine dependence, and recreational illegal drug use (lifetime use <5 occasions each drug) with the exception of occasional cannabis and alcohol use. For both groups, exclusion criteria were clinically significant somatic diseases, head injury or neurological disorders, family history of schizophrenia or bipolar disorder, and use of prescription drugs affecting the CNS. Additional exclusion criteria for both study groups were native tongue other than German, MRI ineligibility due to non-removable ferromagnetic objects or claustrophobia, pregnancy, ager lower than 18 years, or older than 60 years. Participants were asked to abstain from illegal substances for a minimum of three days and from alcohol for at least 24 hours prior to the imaging session. All participants provided written informed-consent in accordance with the Declaration of Helsinki and were compensated for their participation. The study was approved by the local ethic committee of the canton Zurich.

### Clinical Assessment

Drug use was assessed with the Interview for Psychotropic Drug Consumption developed by Quednow et al. (25). The Obsessive Compulsive Cocaine Use Scale (OCCUS) was used to capture long-term cognitive characteristic of craving for cocaine (21). The brief version of the Cocaine Craving Questionnaire (CCQ) was used to measure current cocaine craving (26). The ability to use visual mental imageries was assessed with the Betts Questionnaire Upon Mental Imagery (QMI) (27), the Richardson Controllability Questionnaire (RCQ) (28), the Guy Emotive Imaging Scale (GEIS) (29) and the Spontaneous Use of Imagery Scale (SUIS) (30). Additional clinical and neuropsychological assessments are presented in the Supplementary Methods.

### FMRI Acquisition and Setup

Each participant completed one session in a Philips Achieva 3.0 Tesla magnetic resonance (MR) scanner with an eight channel SENSE head coil (Philips, Best, The Netherlands) at the MR Center of the Psychiatric Hospital, University of Zurich. To identify the VTA/SN using BrainVoyager QX v2.3 (Brain Innovation, Maastricht, The Netherlands), anatomical images were acquired using a spin-echo T2-weighted sequence with 70 sagittal plane slices of 230×184 mm^2^ resulting in 0.57× 0.72× 2 mm^3^ voxel size. Functional data were acquired in 27 ascending transverse plane slices using a gradient-echo T2*-weighted echo planar image sequence with in-plane resolution 2× 2 mm^2^, slice thickness 3 mm, slice gap 1.1 mm, field of view 220× 220 mm^2^, TR/TE 2000/35ms, and flip angle 82°. The slices were aligned with the anterior-posterior commissure. Each participant performed four 7 min fMRI runs (195 volumes). Individual brain volumes were converted from Philips PAR/REC format to ANALYZE DRIN using software from Philips and then placed on a server in real time. The BOLD signal was extracted from the PAR/REC files on a second computer running TurboBrainVoyager (TBV) v3.0 (Brain Innovation, Maastricht, The Netherlands). During the two neurofeedback runs, VTA/SN activity was provided to the participant in the scanner via MR compatible goggles using a custom presentation software developed in Microsoft Visual Studio 2008 (Microsoft, Redmond, WA, USA).

### Neurofeedback Task and Instruction

#### Prescanning Procedure

Outside the scanner, participants were instructed about the goal of the experiment, i.e. to gain self-control over the reward-related brain regions by imagining non-drug related rewarding stimuli. To assess the ability of generating vivid mental imagery, we used an adapted version of the Prospective Imagery Task (PIT) (31, 32): we provided a list of five potentially rewarding sceneries/topics (i.e., positive experiences with family and friends, professional achievements, romantic or sexual memories, hobbies, delicious food including positive scents) plus two individually defined topics, which they rated according to speed (how rapidly mental images can be generated), vividness, and detail on a scale from 1–10. Only the three best ranked topics were used during scanning.

#### Neurofeedback Task

First, each participant underwent an anatomical T2-weighted scan to identify the VTA/SN. The location of this brain region was selected based on previous research (33, 34). The caudal edge of the SN is determined by the cranial edge of the pons at the midline. The cranial border of this region overlaps with the cranial border of the tegmentum. The VTA was determined by the anterior connection between the two lateral SN structures. Both regions were combined into a single region of interest (ROI), which was then coregistered with the functional scans in TBV during the neurofeedback runs. We used the same neurofeedback paradigm as recently published by Sulzer et al (18). The experiment consisted of four runs: a pre-training imagery run, two imagery runs with neurofeedback and a post-training imagery run. Each run comprised nine blocks of alternating “Rest” (20s) and “Happy Time” (20s) conditions. During the “Happy Time” condition, participants were asked to raise the position of the smiley on the screen as high as possible using non-drug rewarding mental imagery. The position and color of the smiley were proportional to the current BOLD signal of the VTA/SN. As the smiley rose, its color gradually changed from red to yellow. During the “Rest” condition, participants were asked to perform a distraction task such as mental arithmetic or imagined paper writing, thereby reducing the height of the smiley and making it redder in color. During the preand post-training imagery runs, the instructions “Happy Time” and “Rest” were provided without smiley feedback.

## Data Analysis

### FMRI ROI Analysis

**Preprocessing**

Data were realigned, slice-timing corrected (35), coregistered for each participant to its individual T2 space and spatially smoothed with a 4 mm full width at half maximum Gaussian kernel using SPM8.

### First and Second Level Analysis

Data analyses were performed in SPM (SPM8, build 6906, http://www.fil.ion.ucl.ac.uk/spm/software/spm8/) using a general linear model (GLM) analysis. In the first level analysis, we specified a GLM with regressors for the “Happy Time” and the “Rest” conditions. The canonical hemodynamic response function in SPM8 was used for convolving all explanatory variables. To test for significant mental imagery induced activity, we contrasted “Happy Time” vs. “Rest” and included the six movement regressors (3 rotations, 3 translations) of the realignment to account for residual motion artefacts. In the second level, we extracted the contrast estimates of the reward imagery contrast (“Happy Time” – “Rest”) from the subject-specific anatomical VTA/SN ROIs. First, we addressed whether CU were able to induce VTA/SN activity with reward imagery performing a one-sample t-tests of the reward imagery contrast across all runs. Second, to test whether obsessive-compulsive thoughts impair the ability to self-regulate the VTA/SN with reward imagery we estimated Spearman correlations between the mean VTA/SN beta estimates (across all four runs) and the OCCUS score as well as lifetime cocaine consumption (in grams). To assess the enhancing effect of NFB, we estimated a paired t-test comparing the mean VTA/SN beta estimates (“Happy Time” vs. “Rest”) during the two NFB training runs with that of the pre- and post-training runs ((mean beta estimates NFB runs 1+2) – (mean beta estimates pre- and post-training run)). Finally, we performed a two-sample t-test between CU and HC to test for any difference in VTA/SN activity and NFB effects. Please note that activity differences between “Happy Time” and “Rest” were not caused by physiological artefacts, as the differences in heart rate and respiration

Self-regulation of the VTA/SN in cocaine users between the two conditions did not correlate with brain activity differences (see Supplementary Methods).

### FMRI Whole-brain Analysis

For image preprocessing and whole brain analysis please see supplemental methods.

### Statistical Notes

Normal distribution was tested with the Kolmogorov-Smirnov test and non-parametric test were used for non-normally distributed data. Huynh-Feldt corrections were utilized to correct for sphericity violations. We applied Bonferroni-corrected pairwise comparisons as post hoc tests for significant main effects. Finally, the correlation analyses were controlled for multiple comparisons using Bonferroni correction.

## Results

### Demographics

The initial study sample comprised 60 participants (CU=30, HC=30). In the CU group, one participant had to be excluded because of opioid dependence, two participants refused to take part in the fMRI experiment, two participants cancelled the scanning due to discomfort, two participants were excluded because of negative cocaine hair analysis and one participant because of signal loss in functional images. Additional, one HC was excluded due to MRI ineligibility (head size), one HC was excluded due to signal loss in functional images. The final sample consisted of 50 participants: 22 CU and 28 HC. The main route of cocaine administration was intranasal in 20 CU, while two CU were primarily inhaling cocaine. Of the 22 CU, 11 fulfilled the criteria of cocaine dependence, three had cocaine substance abuse and eight individuals were recreational users. All participant demographics, clinical data, and group comparisons are summarized in Table 1 and Table S2.

### Behavioral Data

#### Intact Subjective Valuation of Ability to Imagine Rewards in CU

According to the PIT measures, neither symptom severity of obsessive-compulsive thoughts nor the amount of cocaine use impaired the ability to imagine rewarding non-drug related scenes (OCCUS: r=-.207, p=.355; lifetime cocaine consumption: r=-.034; p=.882). Also, compared to HC, CU showed no differences in the ability to imagine rewards (PIT: T=-1.63, p=.11), in the vividness (QMI, GEIS) as well as controllability of mental images (RCQ), and in the tendency to use mental images in daily life (SUIS) (see Table 1). The subjective ability to use mental imagery hence appeared intact in the current sample of CU. A debriefing after the scan confirmed that all CU have used non-drug reward imagery to self-regulate the VTA/SN activity. However, ten CU reported sporadic involuntary thoughts about cocaine during the fMRI scan, predominantly at the end of the experiment (last NFB run and Transfer run).

**Table 1.**
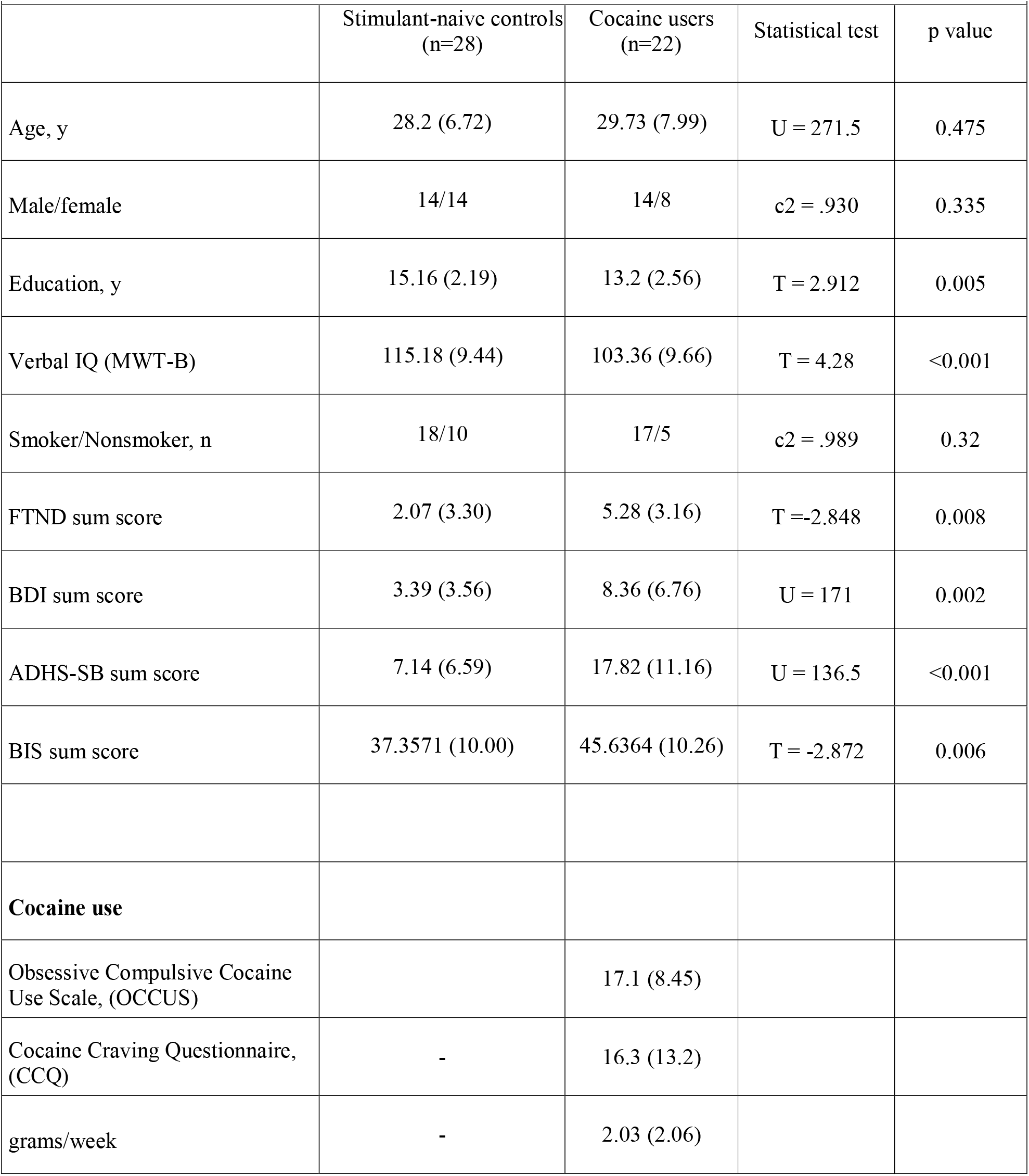
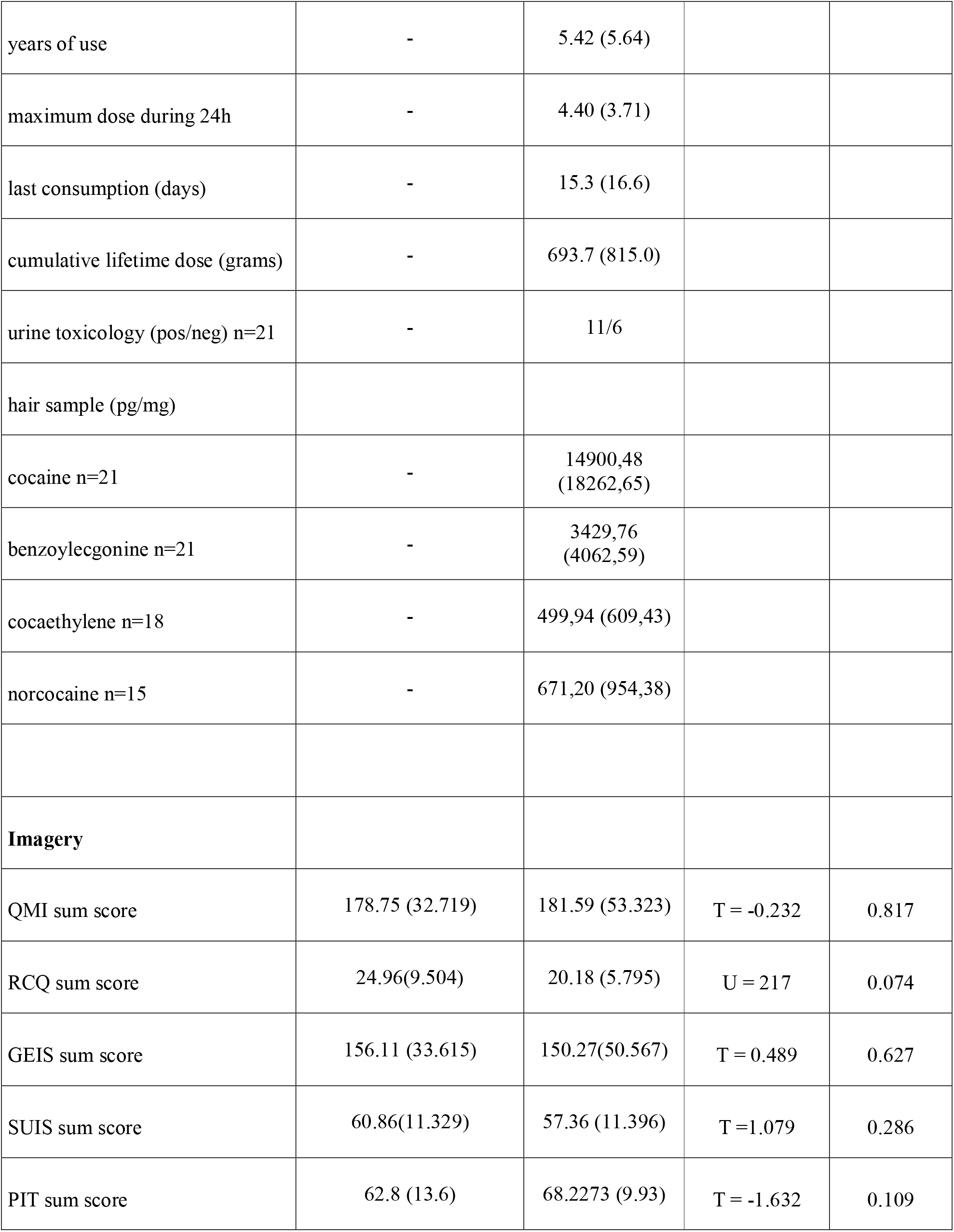
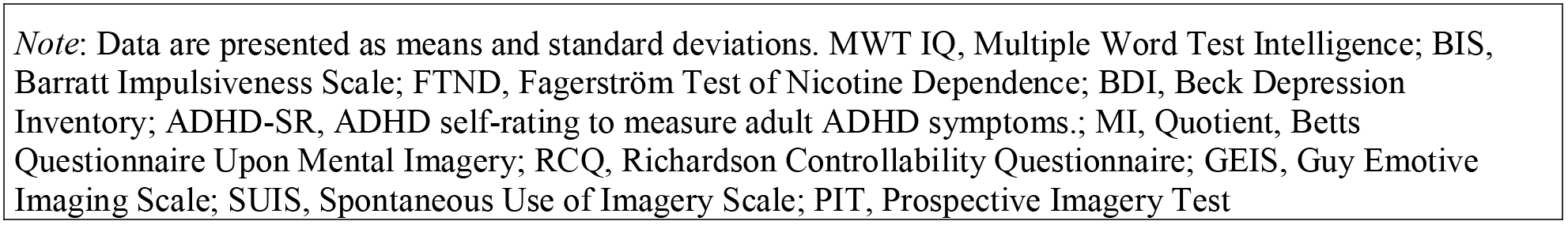
Demographic, clinical data and cocaine use

| Table 1. Demographic, clinical data and cocaine use |  |  |  |  |
| --- | --- | --- | --- | --- |
|  | Stimulant-naïve controls<br>(n=28) | Cocaine users<br>(n=22) | Statistical test | p value |
| Age, y | 28.2 (6.72) | 29.73 (7.99) | U = 271.5 | 0.475 |
| Male/female | 14/14 | 14/8 | c2 = .930 | 0.335 |
| Education, y | 15.16 (2.19) | 13.2 (2.56) | T = 2.912 | 0.005 |
| Verbal IQ (MWT-B) | 115.18 (9.44) | 103.36 (9.66) | T = 4.28 | <0.001 |
| Smoker/Nonsmoker, n | 18/10 | 17/5 | c2 = .989 | 0.32 |
| FTND sum score | 2.07 (3.30) | 5.28 (3.16) | T = -2.848 | 0.008 |
| BDI sum score | 3.39 (3.56) | 8.36 (6.76) | U = 171 | 0.002 |
| ADHS-SB sum score | 7.14 (6.59) | 17.82 (11.16) | U = 136.5 | <0.001 |
| BIS sum score | 37.3571 (10.00) | 45.6364 (10.26) | T = -2.872 | 0.006 |
| **Cocaine use** |  |  |  |  |
| Obsessive Compulsive Cocaine Use Scale, (OCCUS) |  | 17.1 (8.45) |  |  |
| Cocaine Craving Questionnaire, (CCQ) | - | 16.3 (13.2) |  |  |
| grams/week | - | 2.03 (2.06) |  |  |
| years of use | - | 5.42 (5.64) |  |  |
| maximum dose during 24h | - | 4.40 (3.71) |  |  |
| last consumption (days) | - | 15.3 (16.6) |  |  |
| cumulative lifetime dose (grams) | - | 693.7 (815.0) |  |  |
| urine toxicology (pos/neg) n=21 | - | 11/6 |  |  |
| hair sample (pg/mg) |  |  |  |  |
| cocaine n=21 | - | 14900,48<br>(18262,65) |  |  |
| benzoylecgonine n=21 | - | 3429,76<br>(4062,59) |  |  |
| cocaethylene n=18 | - | 499,94 (609,43) |  |  |
| norcocaine n=15 | - | 671,20 (954,38) |  |  |
| **Imagery** |  |  |  |  |
| QMI sum score | 178.75 (32.719) | 181.59 (53.323) | T = -0.232 | 0.817 |
| RCQ sum score | 24.96(9.504) | 20.18 (5.795) | U = 217 | 0.074 |
| GEIS sum score | 156.11 (33.615) | 150.27(50.567) | T = 0.489 | 0.627 |
| SUIS sum score | 60.86(11.329) | 57.36 (11.396) | T =1.079 | 0.286 |
| PIT sum score | 62.8 (13.6) | 68.2273 (9.93) | T = -1.632 | 0.109 |

*Note:* Data are presented as means and standard deviations. MWT IQ, Multiple Word Test Intelligence; BIS, Barratt Impulsiveness Scale; FTND, Fagerström Test of Nicotine Dependence; BDI, Beck Depression Inventory; ADHD-SR, ADHD self-rating to measure adult ADHD symptoms.; MI, Quotient, Betts Questionnaire Upon Mental Imagery; RCQ, Richardson Controllability Questionnaire; GEIS, Guy Emotive Imaging Scale; SUIS, Spontaneous Use of Imagery Scale; PIT, Prospective Imagery Test

## VTA/SN ROI Analyses

### Induction of VTA/SN Activity with Reward Imagery

The first aim of the study was to test whether CU can induce VTA/SN activity with rewarding non-drug imagery. There was a significant difference in BOLD signal between “Happy Time” and “Rest” across all four runs (t=6.176; p<.0001; see Supplementary Results for individual runs). This suggests that CU were able to induce VTA/SN activity by means of rewarding non-drug imagery with and without NFB.

### Reduced VTA/SN Activity Is Associated with Obsessive-compulsive Thoughts and Amount of Cocaine Use

Second, we hypothesized that both obsessive-compulsive thoughts and severity of cocaine use would impair the ability to induce VTA/SN activity with non-drug imagery. We assessed this by correlating OCCUS total scores and lifetime cocaine consumption with the average difference in VTA/SN BOLD signal between “Happy Time” and “Rest” conditions across all four runs. As hypothesized, both correlations were negative (OCCUS total: r_s_=-.495, p<.01, Bonferroni adjusted p=.018; lifetime cocaine consumption r_s_ =-.393, p<.05, Bonferroni adjusted p=.07).

### NFB enhances the Induction of VTA/SN Activity through Mental Imagery

We examined whether NFB enhanced the induction of VTA/SN activity through imagery by comparing the reward imagery contrast between the runs with NFB to those without NFB. This analysis revealed significantly stronger activation during reward imagery in the NFB runs (t=2.777, p<.05). However, although NFB itself was effective, the two NFB runs did not result in a persistent training effect at the end of the imaging session as there was no significant difference between the pre- and post-NFB runs (t=1.810, p=.251).

### Group Comparison of Self-induced VT/SN Activity and NFB between CU and HC

In addition to our primary dimensional analyses within the CU group, we included a HC group to examine group differences. Strikingly, we observed no differences between HC and CU in the overall ability to up-regulate VTA/SN activity across all runs (t=.086, p=.932) nor in the effects of neurofeedback (t=.345, p=.732). Data on the HC group alone are presented in the Supplementary Results).

## Whole-brain Analyses

### Activation of the Reward Network with Mental Imagery

We hypothesized that mental imagery of non-drug rewards would activate regions throughout the reward network within the complete sample (CU + HC). As predicted, the reward imagery contrast revealed strong activation across several regions of the dopaminergic reward system including the VTA/SN complex, ventral (VS) and dorsal striatum (DS), medial prefrontal cortex (mPFC), hippocampus, insula, and posterior cingulate cortex PCC (Figure 4, Table S5) (p<.05 peak-level FWE whole-brain corrected, with a cluster-defining threshold of p<.001 uncorrected).

**Figure 1.**
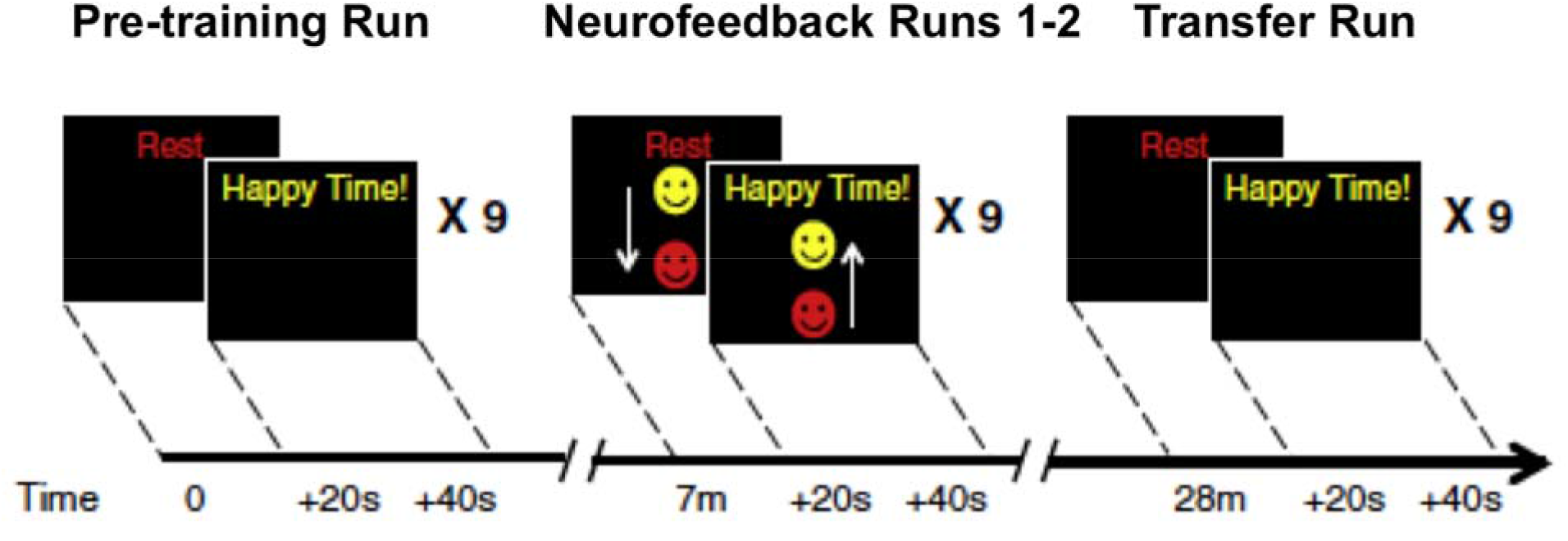
Task Design adapted from the previous publication of Sulzer et al. (18). Following an anatomical localizer, each participant underwent four runs, each one composed of “Rest” (20 s) followed by “Happy Time” (20 s), then repeated nine times. The first and last runs (pre-training and post-trainining) only showed instructions with no visual Neurofeedback. During the two Neurofeedback runs, we instructed participants to use rewarding non-drug imagery to raise the ball during “Happy Time”, and neutral imagery to lower the ball during “Rest”.

**Figure 2.**
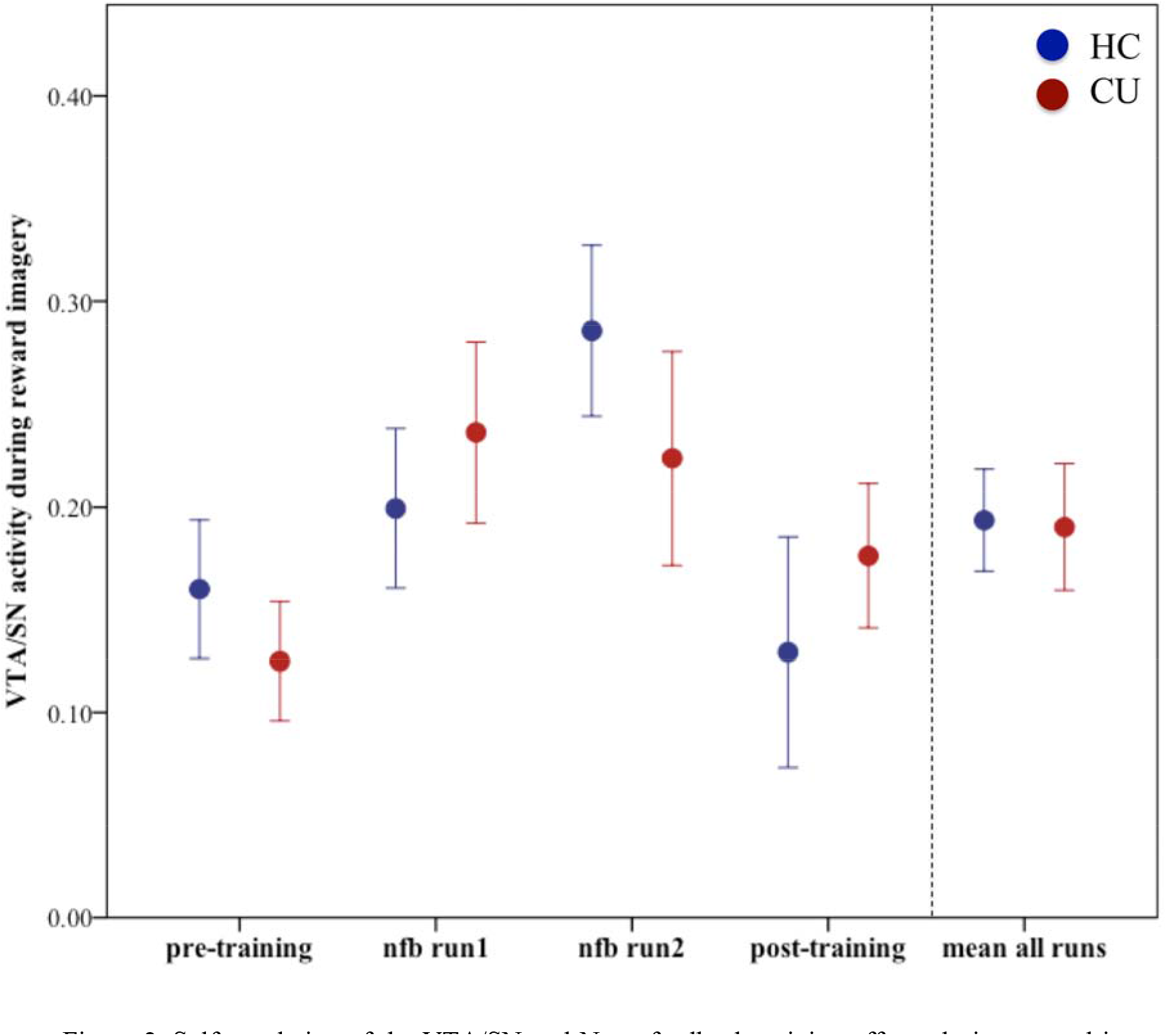
Self-regulation of the VTA/SN and Neurofeedback training effects during reward imagery. The reward imagery contrast estimate (“Happy Time” – “Rest”) is plotted for each run separately and as mean across all runs. Error bars indicate 1 SEM. CU, cocaine users. HC, healthy controls, nfb neurofeedback.

**Figure 3.**
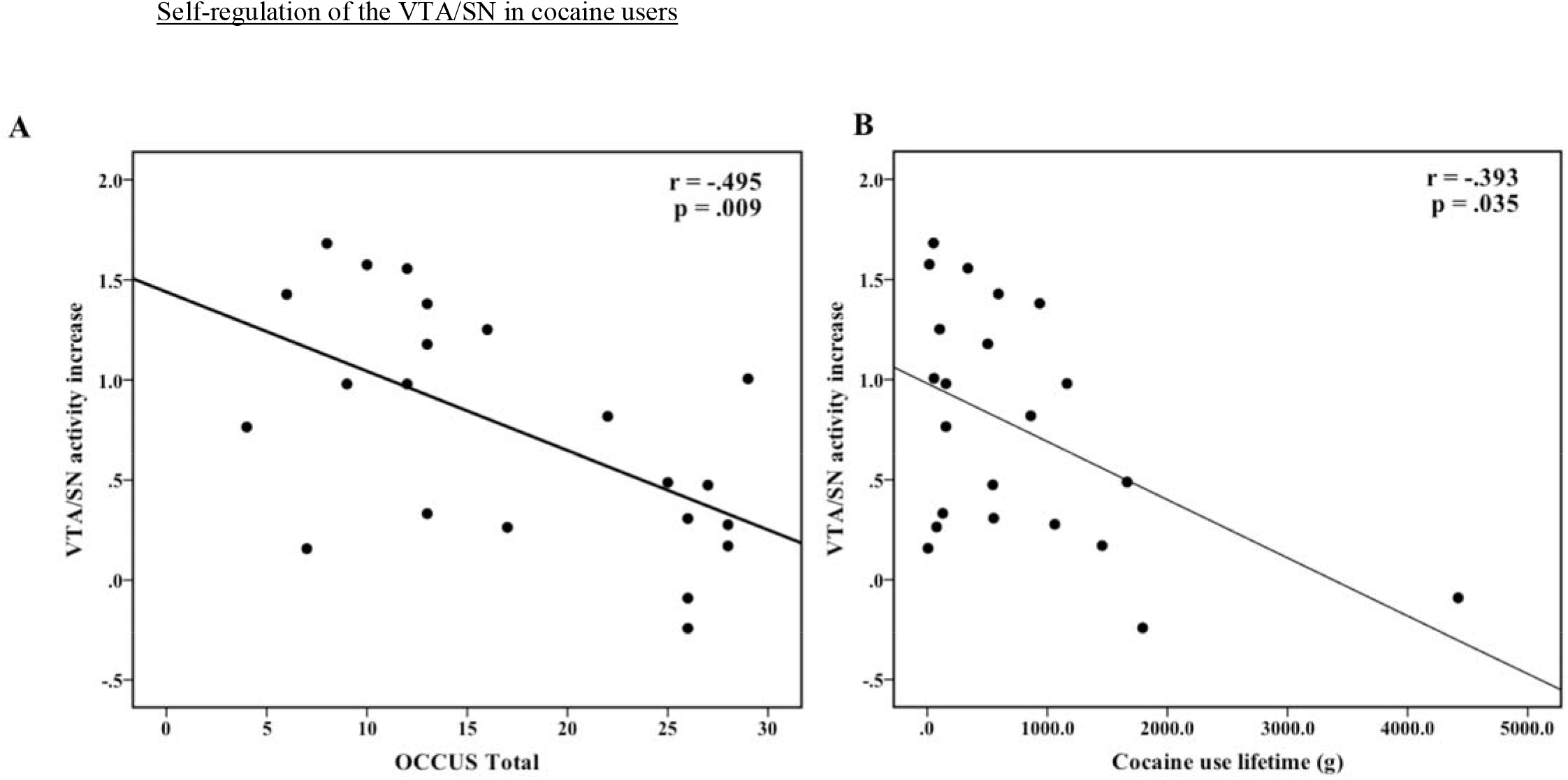
Spearman correlation of the reward imagery contrast estimate (“Happy Time”-”Rest”) with (A) severity of obsessive-compulsive thought about cocaine (OCCUS Total score) and (B) lifetime cocaine consumption (in g).

**Figure 4.**
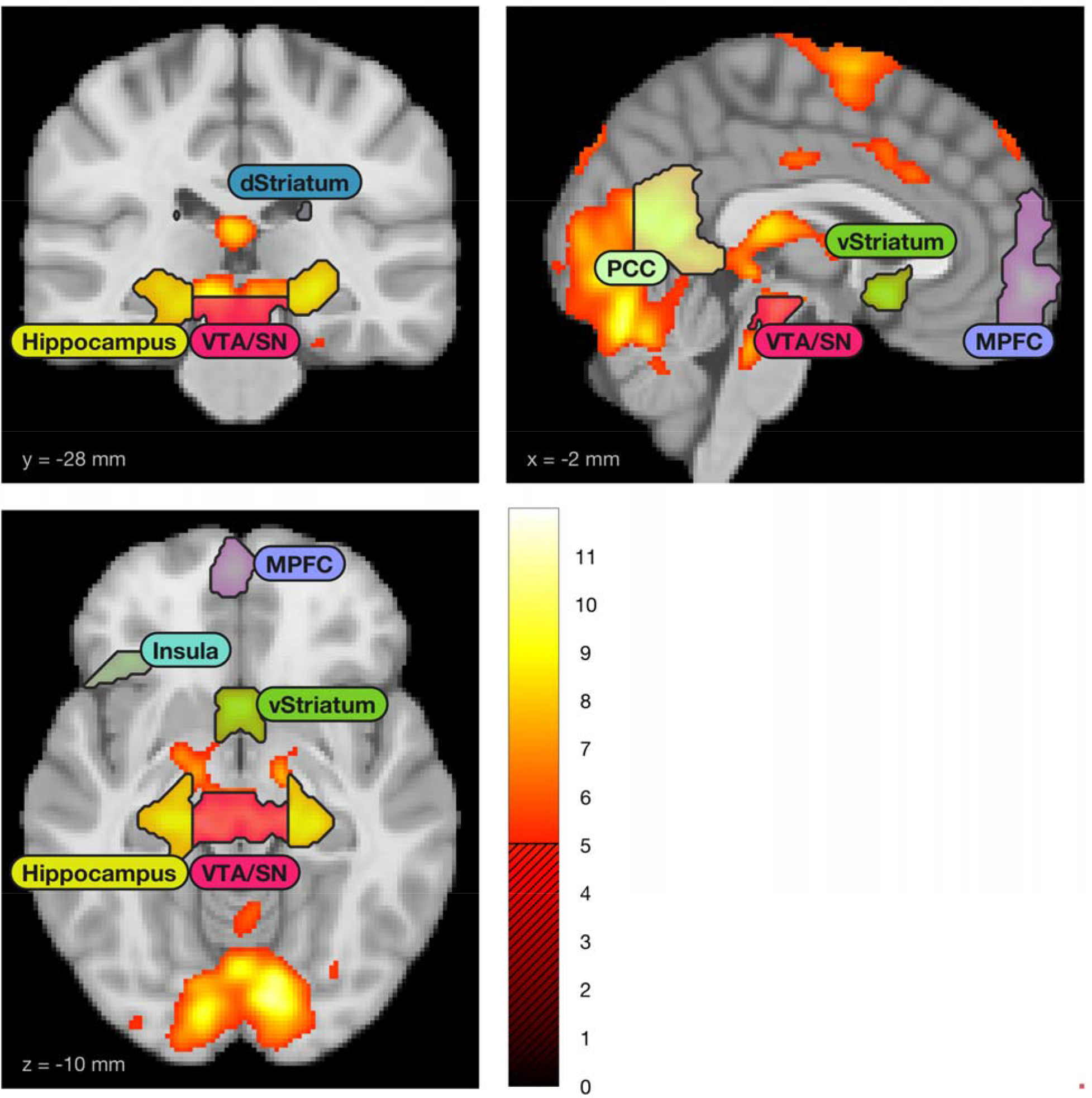
Voxel-wise whole brain analysis of the reward imagery contrast (“Happy Time” – “Rest”) across the complete sample (CU + HC, n=50), peak-level corrected, FWE<.05. Analysis revealed significant activation in the dopaminergic midbrain and throughout the reward network during reward imagery.

## Discussion

Cocaine users’ power to imagine non-drug rewards was behaviorally unimpaired and induced activity in the dopaminergic midbrain and other regions throughout the reward network. The impact of mental imagery was, however, reduced in those with more severe obsessive-compulsive drug use, and in those with higher lifetime consumption. On a group level, we observed no differences between CU and HC. NFB enhanced the effect of non-drug reward imagery, but did not result in transfer effects at the end of our single imaging session. This study represents the first application of reward-based rtfMRI NFB of the dopaminergic midbrain to a clinical population and contributes to the growing field of cognitive neurostimulation as a potential therapeutic approach in psychiatric disorders (19, 36, 37).

### Implications of Impaired Sensitivity to Imagined Rewards in CUD

The association between impaired self-regulation of the VTA/SN by using non-drug related imagery and severity of obsessive-compulsive drug use suggests an important link between maladaptive cognitive features of CUD and dysfunction in reward processing at the neural level. In contrast, at the behavioral level, neither obsessive-compulsive thoughts nor lifetime consumption was associated with the ability to generate vividly images of non-drug-related rewards. Furthermore, subjective ratings of the ability to imagine rewards did not predict the neural response to imagined rewards (PIT, r_s_=-.081, p=-.719, see Supplementary Results). In other words, symptom severity of CUD was directly linked with impaired neural reward sensitivity and was not related to the individual capability to imagine non-drug rewards. This dissociation between intact subjectively reported reward imagery and impaired neural response in reward circuits provides one possible mechanism for the failure to engage in adaptive goal-directed behavior in CU. Indeed, intact neural activity during reward imagery is relevant for decision-making. For instance, neural responses to imagined rewards reduce the temporal discounting of future rewards and guide choice behavior (38, 39). Translating these findings to the maladapted reward sensitivity in CUD suggests that training imagery of non-drug related rewards may help individuals with CUD to reduce impulsive drug seeking in favor of functional non-drug related decision making. Clinical interventions such as cognitive behavioral therapy and motivational enhancement therapy already aim to improve the intrinsic motivation for adaptive goal-directed behavior (40–43). In conjunction with these psychosocial interventions non-drug related reward imagery and self-activation of the reward circuitry (44) may provide an additional tool to directly target impaired reward sensitivity in CUD.

### Relevance of Chronic Craving for Impaired Non-drug Related Reward Sensitivity

Obsessive-compulsive thoughts are a signature of chronic craving, and they were negatively related to VTA/SN activity during reward imagery. Hence, chronic craving may interfere with the sensitivity to non-drug rewards. This is in contrast to measures of acute craving, which did not explain the reward circuitry impairment during non-drug related reward imagery (CCQ, r_s_=.263, p=.238, see Supplementary Results). Given that acute craving is strongly associated with drug-cue reward sensitivity, these divergent findings suggest that chronic and acute craving may be differentiable on a neural level. One caveat is that we did not directly induce craving in our study and hence likely have low power to detect effects related to acute craving. Given that craving is a multidimensional construct spanning conditioning, cognitive and neurobiological components and occurs in different disease states (21, 45), future research should try to directly address these different aspects and disentangle the neural correlates underlying acute craving and chronic obsessive-compulsive thoughts.

### Potential Effects of Cognitive Neurostimulation with NFB in CUD

Previous studies on self-regulation of the dopaminergic midbrain (18, 19) and other studies using recall of rewarding memories to control neural activity (36, 46) support the effectiveness of NFB for non-invasive cognitive neurostimulation in mental disorders (19, 36, 37, 47–51). In line with previous studies, we showed that imagery of non-drug rewards efficiently stimulated reward-related circuitry in CU across a reward network spanning mesolimbic, mesocortical and hippocampal circuits (18, 52, 53). More importantly, the same regions underlie dysfunctional reward sensitization during the development of addictive behavior (7, 54). We therefore speculate that non-invasive cognitive neurostimulation as shown in our study might target the underlying neural correlates of addictive behavior. Of course, substantial limitations still have to be overcome. First, we did not observe a training effect, at least after one single session. This could potentially be addressed through more extensive training. Second, we have not investigated any generalization effect, or indeed any real-life impact on relevant measures. Obvious candidates for relevant clinical outcomes in future training trials would be the impact on cue-induced craving and compulsive drug intake. Indeed, recent rtfMRI NFB studies in depression suggest that even short-term interventions with NFB have lasting impact improving symptom severity and enhancing previous learned cognitive strategies (36, 55).

### Limitations and Open Questions

Recent studies revealed inconclusive findings regarding the generalization and transfer of NFB training when comparing pre- and post-training VTA/SN activity (18, 19). Whereas MacInness et al. found no effect during the pre-training run, but a significant pre- to post-training effect, Sulzer et al. and we found significant pre-training VTA/SN activity, but no significant differences between preand post-training (18, 19). Although speculative, these divergent findings might be explained by differences in task instructions. In the study from MacInness et al (19) the best strategy was explicitly used during the post-training run, which was not the case in our study and the previous one from Sulzer and colleagues (18). Furthermore, in our study, participants underwent a pre-scanning training, which might have improved the self-regulation ability in the first pre-training run. The lack of NFB transfer effects might also be because our training was limited to one single scan session. This might have caused fatigue and adaptation of the dopamine signal especially during the last post-training run, thus obscured potential transfer effects. Future NFB studies should use longitudinal designs with multiple training sessions to identify potentially lasting transfer effects. Longitudinal designs will also allow for assessing real-life impact on relevant clinical measures.

Real-time fMRI NFB is a complex and expensive intervention that will face substantial cost-effectiveness hurdles. However, given the chronic nature of CUD and the limited treatment options with no approved pharmacological interventions, it is imperative to pursue all novel treatment options. Also, there is accumulating evidence that only a few NFB training sessions produce effects that last for several months up to a year (56, 57). Other advantages are that NFB is safe (58) can be personalized, combines psychological (i.e. mental strategies) as well as biological (i.e. brain changes) factors, and focuses on learning to self-heal (36, 55).

Finally, our broad study sample included a wide range of CU from recreational to chronically compulsive drug taking. This allowed a dimensional approach to investigate the association between symptom severity and VTA/SN self-regulation, but it likely limited our power for detecting categorical differences, which are potentially more pronounced for severe CUD. As this group is of particular clinical relevance, future studies should focus on severe chronic individuals with CUD.

### Conclusion

Cocaine users can voluntary induce dopaminergic midbrain activity by means of non-drug rewarding imagery and improve this ability with rtfMRI NFB. Combining reward imagery and rtFMRI NFB has great potential to modify the imbalance of reward sensitivity and reinstate non-drug reward responsiveness. This motivates further work to examine the potential of cognitive neurostimulation approaches in the treatment of CUD.

## Acknowledgments

We thank Ged Ridgway for his helpful advice. Furthermore, we would like to thank all participants for their time and interest in our study.

